# Prospects of using Avocado oil for attenuating quorum sensing regulated virulence, bio-filming formation and its antibacterial and antioxidant activities

**DOI:** 10.1101/486928

**Authors:** Hanan M. Al-Yousef, Musarat Amina, Syed Rizwan Ahamad, Wafaa H. B. Hassan

## Abstract

Quorum sensing inhibition (QSI) is considered as an attractive strategy for the development of anti-pathogenic agents, mainly for drug resistant bacteria. The anti-quorum sensing activity was investigated by biosensor bioassay using *Chromobacterium violaceum* CVO26 and *Pseudomonas aeruginosa* PAO1. Quorum sensing is a key regulator of virulence factors of *Pseudomonas aeruginosa* such as bio-film formation, motility, productions of proteases, hemolysin, and Pyocyanin production. Additionally, the GC/MS technique was employed to detect the essential components of avocado oil. Avocado oil inhibits quorum system-mediated virulence factor production such as violacein in C. Violaceum CVO26 and elastase, Pyocyanin production in *Pseudomonas aeruginosa* PAO1. Additionally, the use of sub-minimum inhibitory concentrations (sub-MICs) of avocado oil significantly inhibits the quorum system-mediated biofilm formation, exopolysaccride production (EPS) and swarming motility. Furthermore, this study concerned the potent activity of avocado oil antibacterial and antioxidant agent. Moreover, a total of 23 components was identified in avocado oil by GC/MS.Avocado oil could be exploited as a natural source of anti-pathogenic, where the pathogenicity is mediated through quorum sensing, antibacterial as well as antioxidant agents.

**Graphical Abstract:** 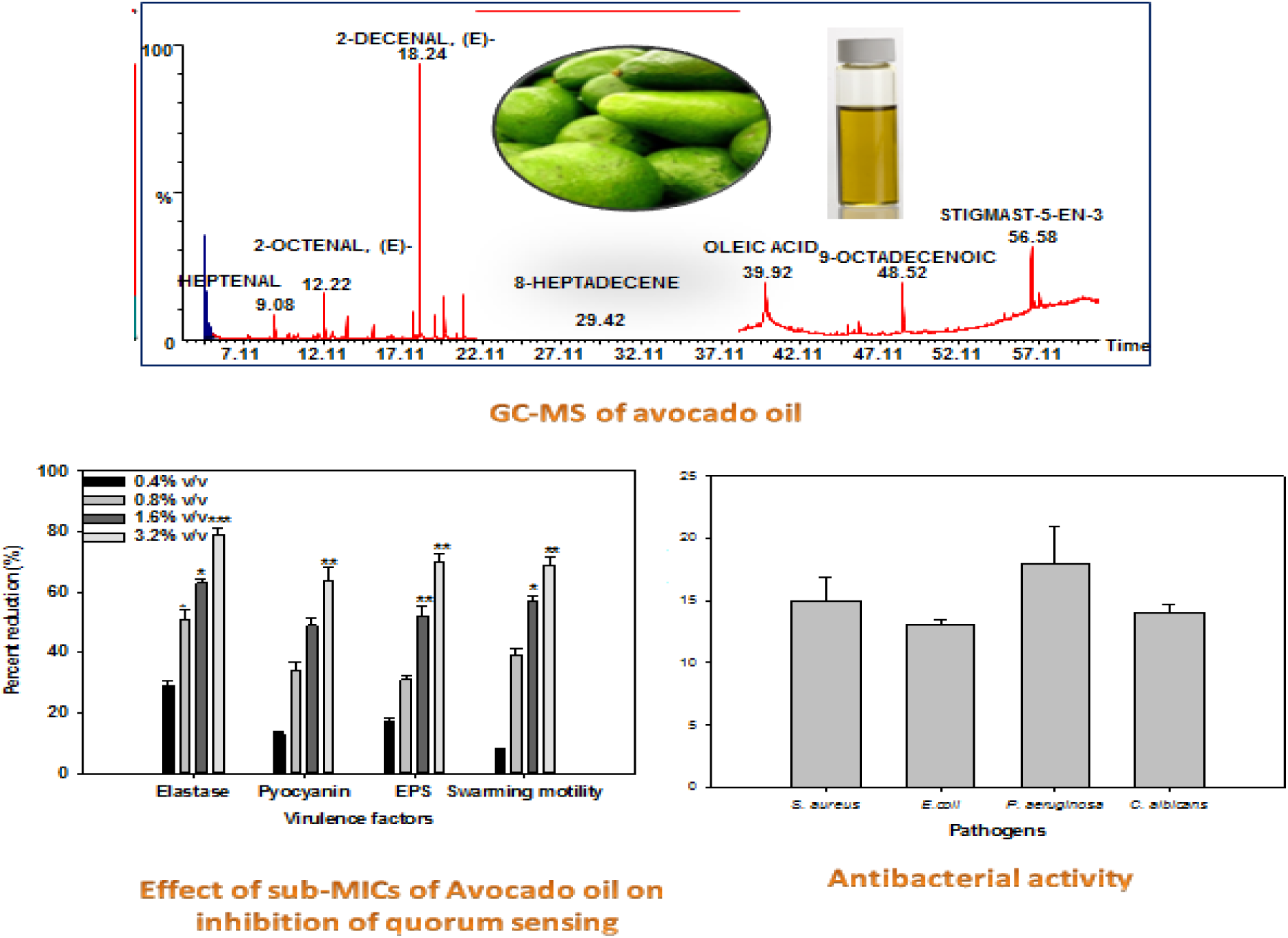

## INTRODUCTION

The *Persea Americana* mill F. (*Avocado*) belonging to the family Lauraceae, commonly known as “*ahuácatl*” (a Mexican word) meaning “testicle” which refers to the shape of the fruits. It is comprised of aromatic shrubs and trees. Avocados have been cultivated for their highly nutritious fruits since about 8,000 BC, and there is evidence that they were eaten as a wild fruit before then (Samson, 1986). The avocado is unique fruit due to variation on its chemical composition, these compositions obviously different according to the time of the seasons, cultivar, soil, environment, etc. The chemical composition of the edible portion of the flesh is water 65-80%; protein l-4%; sugar about 1%; oil 3-30%. It is high in B vitamin and moderately in vitamins A and D. The better recognition about this variation of fruit compositions is important. The avocado oil is considered as high digestible owing to its high oil content; it possesses highest energy value than any other food (Purseglove, 1968). Avocado though highly nutritious fruit yet low in sugar content; therefore, it can be recommended as high energy food for the diabetic (Samson, 1986; Swisher,1988).

Most medical applications of plants magnified on their antimicrobial effect with ignore attention towards anti-pathogenic effects (Wallace, 2004). Nowadays, research efforts are concentrated on controlling microbial infection through developing antipathogenic agents which manage microbial diseases by inhibiting microbial communication process called microbial quorum sensing (QS). It is well known that many pathogenic bacteria used a QS system to regulate genes required for virulence expression; therefore, the inhibition of QS system is obeyed as a new strategy to control the pathogenicity and for the development of anti-pathogenic agents. The release of *Pseudomonas* virulence factors is regulated by a quorum communication system (Zhang and Dong, 2004). Quorum sensing in *P. aeruginosa* is regulated by signaling molecules claimed N-acylated homoserine lactones (AHLs). The concentration of these molecules arises in relation to the high of the bacterial population, those signaling molecules return to the bacteria to control bacterial pathogenicity (Fuqua and Greenberg, 2002). Thereby, removal of QS represents a potential advance system to manage bacterial virulence and resistance (Hong, et al., 2012). Literature studies claim that medicinal plants are the rich source of quorum scavenging compounds (Mohamed, et al., 2014; Koh and Tham, 2011; Choo, et al., 2006). So, this study assessed the quorum sensing inhibition (QSI) effect of avocado oil using the reporter *Chromobacterium violaceum*. This oil showed QSI activity was investigated for anti-pathogenic potential against *Pseudomonas aeruginosa* PAO1. In this respect, their influence on the virulence of *P. aeruginosa* was examined, including biofilm formation (BF).

The literature survey revealed that there is no such reported data about commercial avocado oil. This prompted us to investigate the oil aiming to identify its chemical constituents by Gas chromatography (GC) and gas chromatography mass spectrometry (GC/MS) and compare it with the reported data. In addition to be explored for their QSI properties. In this respect, the chemical compounds of the extra virgin oil were detected by using GC/MS analysis. Considering the various medicinal and functional properties of avocado oil, a study was also planned with the aim to determine the QS and biofilm inhibitory properties (BI) of this oil against pathogenic bacteria. Moreover, it may need further work to evaluate its different biological and pharmacological activities.

## RESULTS AND DISCUSSION

### RESULT

Avocado oil under investigation is dark yellow in color with characteristic odor, soluble in ether and chloroform, insoluble in water. Analysis of avocado oil by GC and GC/MS resulted in the identification of 23 compounds representing 98.67% of the total oil, Figures 1 and 2, their retention indices and area percentages (concentrations) are summarized in Table 1. Antimicrobial screening of essential oil of avocado was determined against *S. aureus, E. coli, P. aeruginosa and C. albicans* 100 µl of avocado oil at a dose of 100 mg/ml test by determination of the zone of inhibition and showed high inhibit activity against *P. aeruginosa*, and moderate inhibit the activities against *S.aureus*, *C. albicans* and *E. coli*, when compared with control, with MIC values between 1.6-6.4 mg\ml for components of avocado oil, Figure 3 and Table 2. The antioxidant activity of avocado oil was evaluated by using DPPH. DPPH-radical scavenging assay showed a significant antioxidant activity (*p* ≤0.05) at a dose dependent matter of avocado oil at different doses (12.5-400 µg/ml) as presented in Table 3, when compare to ascorbic acid and Butylated hydroxyl toluene (BHT), (Figure 4). Quantitative assessment of violacein inhibition in CVO26 by sub-MICs of Avocado oil was shown in Figure 5. A maximum significant inhibition (*p* ≤0.001) of violacein was determined at doses of 0.4 and 0.8 (%v/v) by 80% and 94%, respectively, while at lower doses (0.1 and 0.2 %v/v) a significant reduction in violacein were noticed by 45% (*p* ≤0.05) and 60% (*p* ≤0.005) respectively. So, in the current study sub-MIC concentrations (0.1-0.8 %v/v) of avocado oil were used for further assays (Figure 6a).

**Table 1:** Chemical composition of avocado oil

| S No | Chemical Name | RI | RT | Area% | N Area% |
| --- | --- | --- | --- | --- | --- |
| 1. | Octane | 800.6 | 4.7 | 28.86 | 100 |
| 2. | Heptanal | 901.6 | 7.42 | 0.64 | 2.22 |
| 3. | 2-Heptenal | 956.0 | 9.08 | 2.080 | 7.19 |
| 4. | Octanal | 1002.3 | 10.5 | 0.5 | 1.75 |
| 5. | 3,5-Octadien-2-ol | 1037.6 | 11.58 | 0.69 | 2.4 |
| 6. | Octenal | 1046.2 | 11.85 | 0.64 | 2.2 |
| 7. | 2-Octenal | 1058.1 | 12.22 | 2.19 | 7.59 |
| 8. | 1-octanol | 1074.1 | 12.7 | 0.77 | 2.6 |
| 9. | Nonanal | 1103.7 | 15.2 | 0.93 | 3.24 |
| 10. | Trans-2-Undecanal | 1248.0 | 17.8 | 2.7 | 9.4 |
| 11. | 2-Decenal | 1263.2 | 18.24 | 21.3 | 74.0 |
| 12. | 2,4-Decadienal | 1295.2 | 19.1 | 5.5 | 19.1 |
| 13. | 2,4-Decadienal | 1319.6 | 19.7 | 9.0 | 31.2 |
| 14. | Trans-2-undecanal | 1364.6 | 20.9 | 2.7 | 9.4 |
| 15. | Pentadecane | 1497.9 | 24.34 | 0.36 | 1.25 |
| 16. | 8-Heptadecane | 1674.9 | 29.4 | 0.56 | 1.93 |
| 17. | Hexadecanoic acid | 1979.2 | 36.44 | 0.43 | 1.5 |
| 18. | Oleic acid | 2161.0 | 39.9 | 8.5 | 29.5 |
| 19. | 9-Octadecanoic acid-ester | 2470.0 | 45.1 | 0.45 | 1.56 |
| 20. | Hexadecanoic acid ester | 2515 | 45.8 | 0.77 | 2.6 |
| 21. | 9-Octadecanoic acid ester | 2697.1 | 48.5 | 3.73 | 12.9 |
| 22. | Ergost-5-en-3-ol | 3212.3 | 55.48 | 2.9 | 10.09 |
| 23. | Stigmasitosterol | 3296.0 | 56.58 | 2.9 | 10.09 |

**Table 2.** Minimum inhibitory concentration (MIC) of avocado oil against *S. aureus*, *E. Coli*, *P. aeruginosa* and *C. albicans* bacteria

| Microorganisms | MIC of avocado oil<br>mg/mL | Inhibitory<br>zone (mm) | Control |
| --- | --- | --- | --- |
| *Staphylococcus aureus* (ATCC25922) | 6.4 | 15±0.02 | Ampicillin |
| *Escherichia coli* (ATCC25923) | 3.2 | 14±0.01 | Doxycycline |
| *Pseudomonas aeruginosa* (ATCCPAO1) | 3.2 | 18±0.12 | Doxycycline |
| *Candida albicans* (SC315) | 6.1 | 13±0.23 | Nystatin |

**Table 3:** Free radical scavenging activity of avocado volatile oil

| Name of oil | Percent decolorization by DPPH method |  |  |  |  |  |
| --- | --- | --- | --- | --- | --- | --- |
|  | Concentrations of avocado oil (µg/mL) |  |  |  |  |  |
|  | 12.5 | 25 | 50 | 100 | 200 | 400 |
| Avocado oil | 8.39±1.89 | 31.66±2.45 | 49.27±1.88 | 61.37±5.46 | 70.95±2.52 | 71.85±0.39 |
| Ascorbic acid | 34.91±1.98 | 69.56±2.64 | 87.41±1.47 | 91.5±2.16 | 94.66±2.18 | 95.20±1.65 |

**Figure 1:**
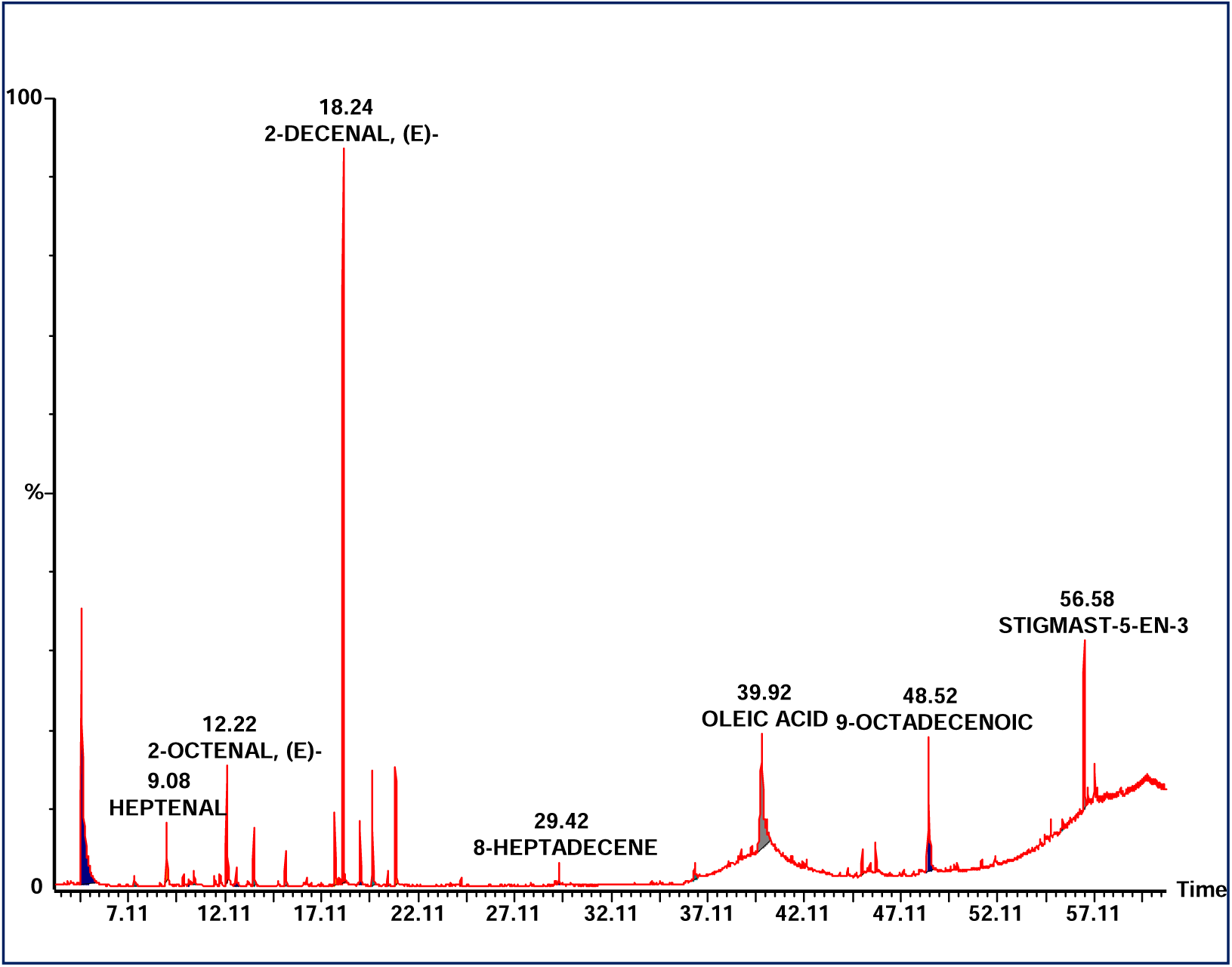
GC/MS chromatogram of the Avocado oil

**Figure 2:**
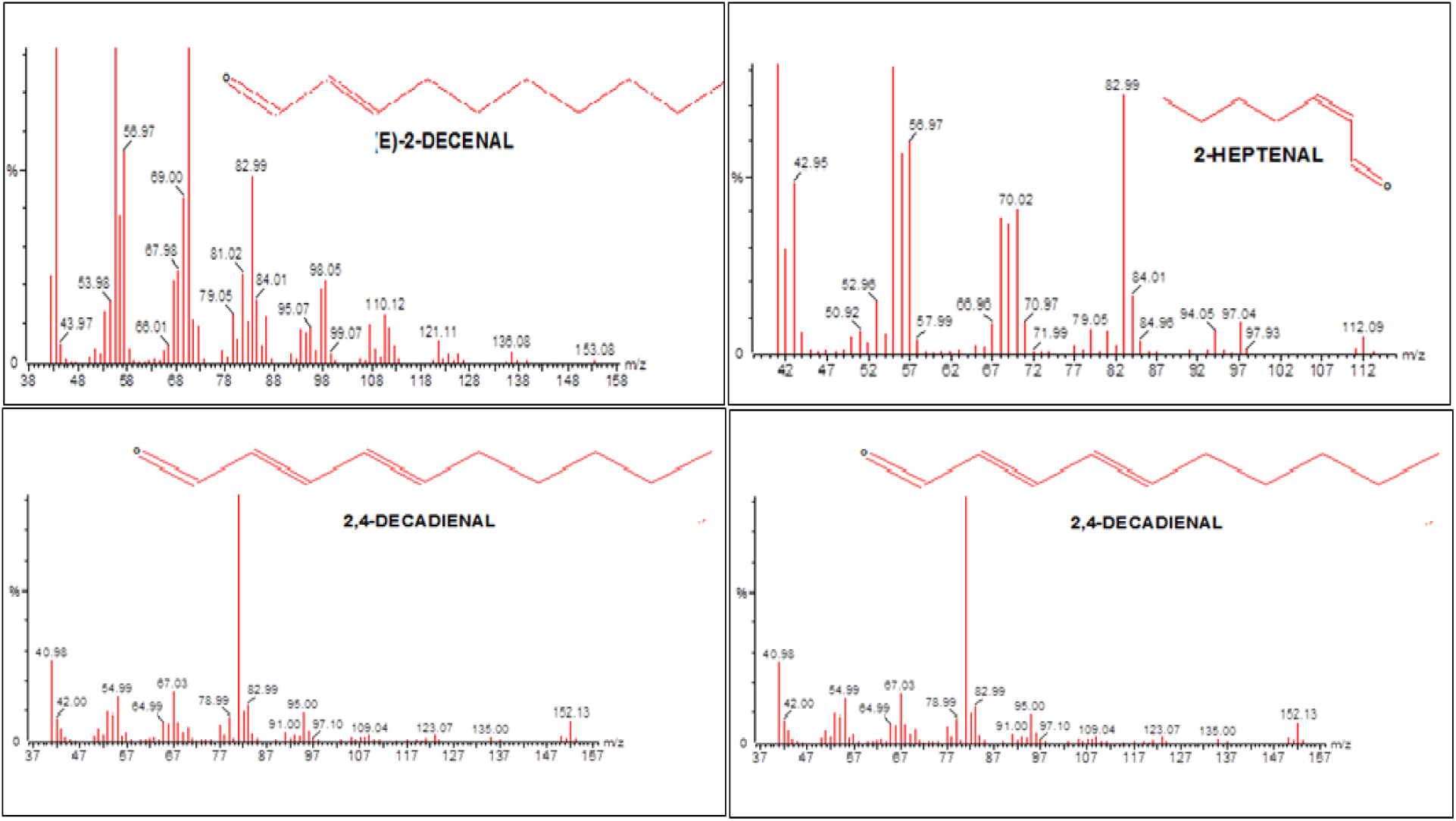
Mass spectra of some essential oil components of avocado oil

**Figure 3:**
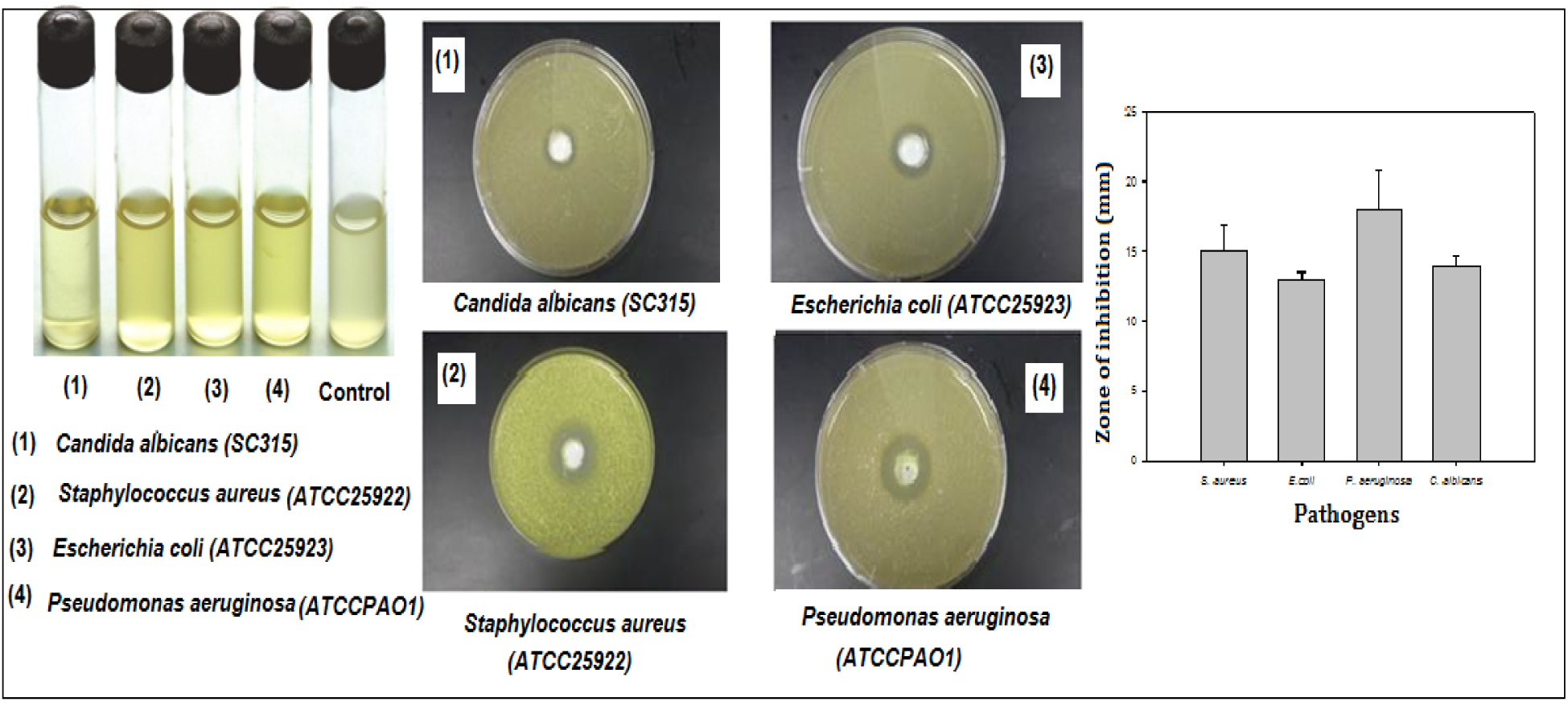
Antimicrobial activity of Avocado oil against different pathogens

**Figure 4:**
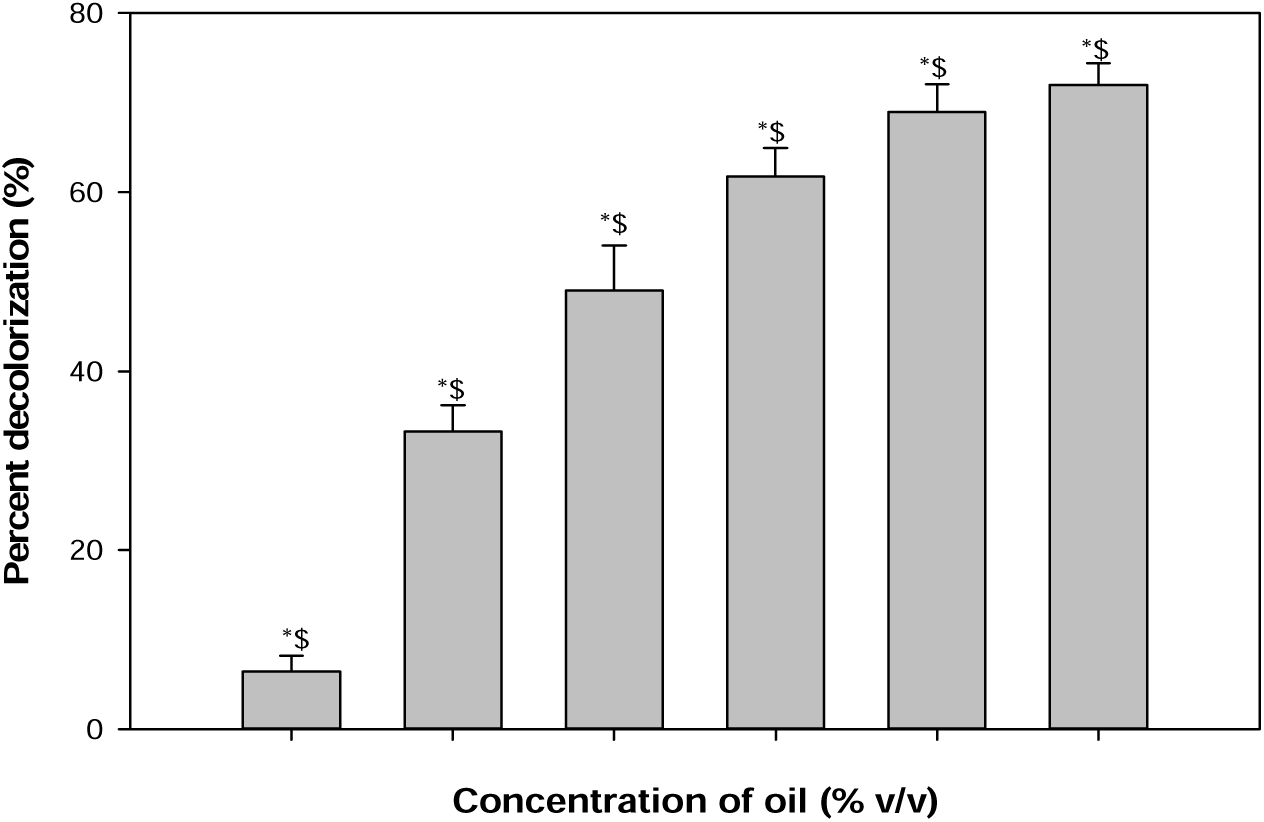
Free radical scavenging activity of Avocado oil by DPPH method. The above data are the mean of three replicates; * shows significant with Ascorbic acid, (p ≤0.05); $ shows significant with Butylated hydroxyl toluene (BHT), (p ≤0.05).

**Figure 5:**
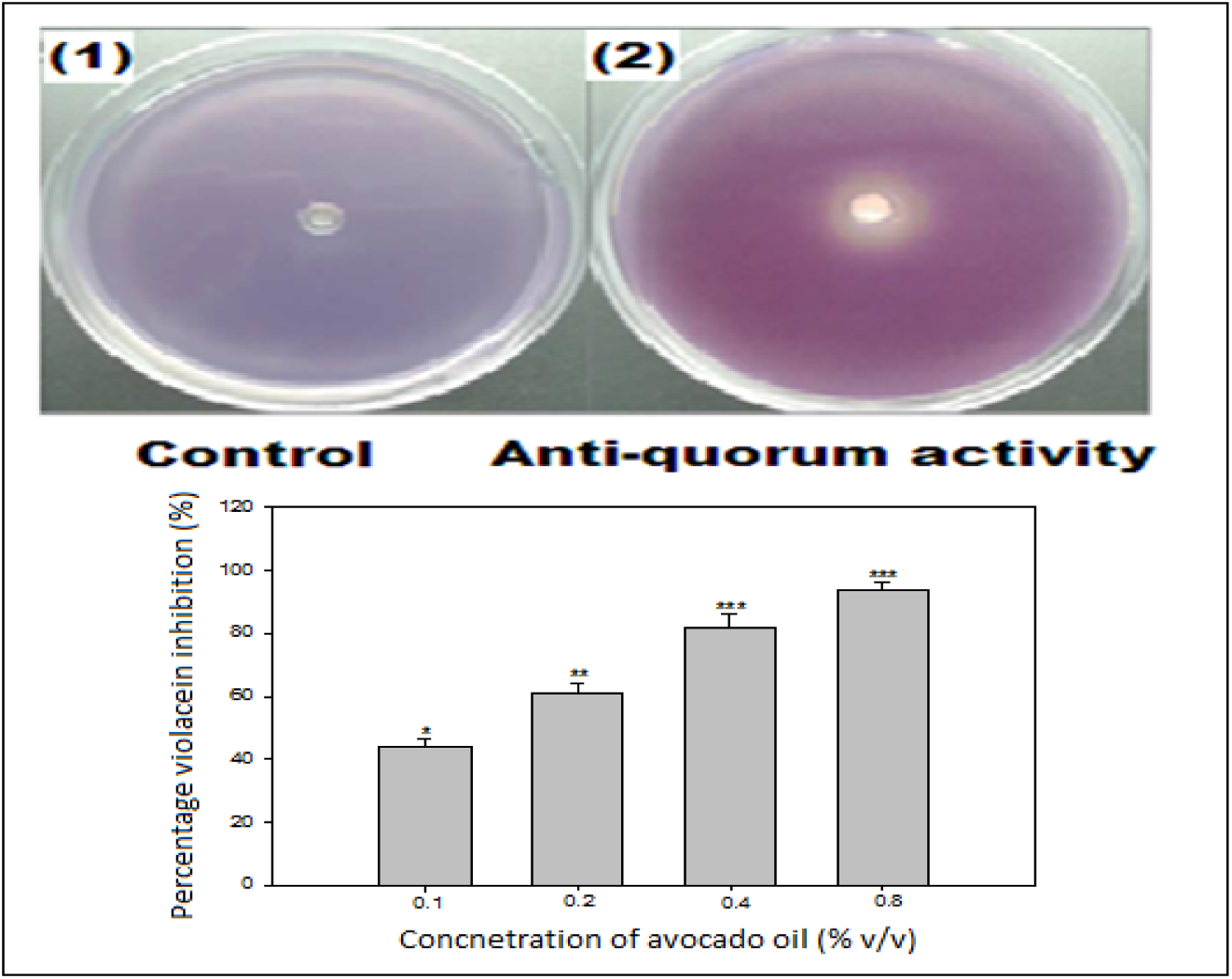
Quantitative assessment of violacein inhibition in CVO26 by sub-MICs of Avocado oil. All of the data are presented as mean ± SD. *, significance at p ≤0.05, **, significance at p ≤0.005, ***significance at p ≤0.00

**Figure 6:**
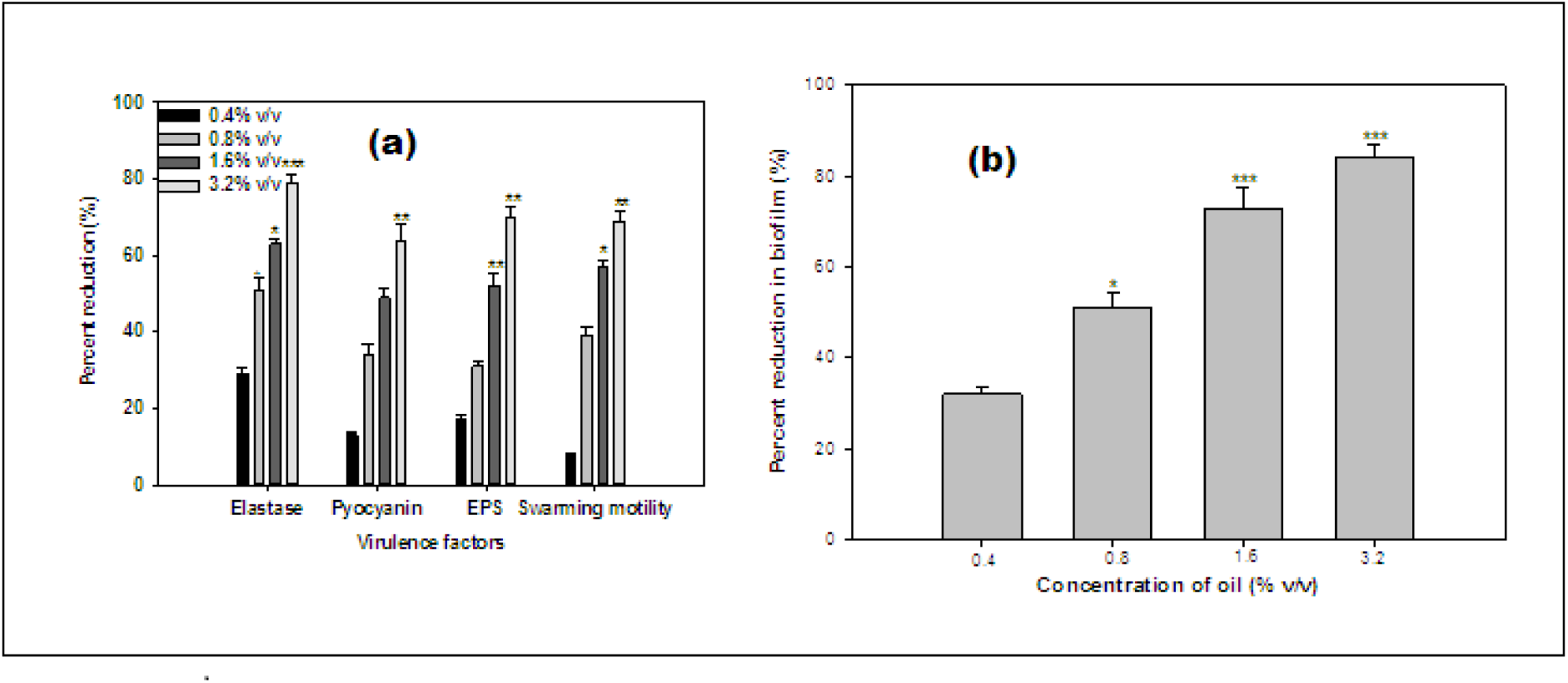
(a) Effect of sub-MICs of Avocado oil on inhibition of quorum sensing regulated virulence factors in *P. aeruginosa* PAO1. All of the data are presented as mean ± SD. *, significance at p ≤0.05, **, significance at p ≤0.005, ***significance at p ≤0.001 and (b) Effect of sub-MICs of Avocado oil on biofilm formation in *P. aeruginosa* PAO1. All of the data are presented as mean ± SD. *, significance at p ≤0.05, **, significance at p ≤0.005, ***significance at p ≤0.001

Statistically significant decrease in **LasB elastase** activity was observed in the culture supernatant of PAO1 treated with sub-MICs of avocado oil. A minimum of 30% inhibition was observed when PAO1 was cultured with avocado oil at a concentration of 0.4 %v/v followed by 52% and 63% (*p* ≤0.05) at doses 0.8 and 1.6 %v/v respectively, moreover, the maximum of 80% (*p* ≤0.001) inhibition was observed at 3.2%v/v concentration of the extract. **Pyocyanin production** (PP) is an important VF produced under QS regulation. The major role of PP is reported in pathogenesis mainly in cystic fibrosis (Winstanley, 2009). A similar reduction in PP was documented in extracts of *Terminalia chebula* (Sarabhai, et al., 2013). A “Pyocyanin” has a green color produced by *P. aeruginosa* (PAOI) after 24-48 h of growth. The disappearance of this pigment indicated the lower levels of PP or no PP is found in the supernatant. The effect of avocado oil on the PP was performed. In the current study, PP level in *Pseudomonas* culture handled with this oil was significantly reduced without affecting the growth of bacteria against the green color of untreated cultures. This might be interpreted as quorum-control of PP (Dietrich, et al., 2006).^[13]^ Moreover, quorum quenching agents have greater impact on PP from *P. aeruginosa* (El-Mowafy et al., 2014; Morkunas et al., 2012). The avocado oil at sub-lethal doses possessed considerable decrease in the PP by PAOI. The maximum significant reduction of 62% (*p* ≤0.005) in PP was recorded at a highest tested concentration (3.2%v/v) followed by 51%, 38%, and 15% in 1.6, 0.8, and 0.4 %v/v concentrations, respectively (Figure 6b).

In the current study, treatment of PAO1 with sub-MICs of avocado oil showed significantly decrement of **exopolysaccharide production** (EPS), the volatile oil concentrations (0.4–3.2 %v/v) demonstrated inhibition in exopolysaccharide production to the level of 19–72%. The maximum significant reduction of 72% (*p* ≤0.005) in EPS was recorded at a highest tested concentration (3.2%v/v) followed by 52% (*p* ≤0.005) at a dose of 1.6 %v/v concentrations, Figure 6a. Similarly, **swarming** migration of PAO1 was also impaired considerably (10–69%) after treatment with avocado oil (0.4-3.2%v/v) concentrations. The maximum significant reduction of 69% (*p* ≤0.005) in swarming was recorded at a highest tested concentration (3.2%v/v) followed by 59% (*p* ≤0.05) at 1.6 %v/v, (Figure 6a). The avocado oil was tested for *Pseudomonas* BF using tube assay method. It showed significant effect on **BF** by *P. aeruginosa* PAOI against control (Figure 6b). It highly inhibited the biofilm biomass in a dose-dependent manner without affecting the *P. aeruginosa* (PAO1). The avocado oil showed 32, 52 (*p* ≤0.05), 72 (*p* ≤0.001), and 83% (*p* ≤0.001) decrease in the BF ability of PAO1 at 0.4, 0.8, 1.6, and 3.2 %v/v of oil concentrations, respectively, (Figure 6b).

## DISCUSSION

Alkanals and hydrocarbons are major composition present in GC/MS analysis. Alcohols are fewer in number than those found by Yamaguchi et al. (1983) in their work on avocado volatiles. As expected, lipid breakdown volatiles, e.g. hexanal, heptanal and decadienal, are clearly evident in the oil. Among these as the main components are octane (**1**, 28.86%), 2-decenal (**11**, 21.3%), 2,4 decadienal (**13**, 9.0%), oleic acid (**18**, 8.5%), 9-octadecanoic acid ester (**21**, 3.73%), ergost-5-en-3-ol (**22**, 2.9%) and stigmasterol (**23**, 2.9%). While pentadecane (**15**, 0.36%), hexadecenoic acid (**17**, 0.43%), and octanal (**4**, 0.5%). occurs as minor constituents of the oil, (Figure 1). By comparison of our results in this study with the previous reports on other avocado oil sources growing in different countries it has disclosed that the chemical composition of avocado oil were completely different from Babol, Iran avocado oil results (Azizi and Najafzadeh, 2008), as well as the results reported in different literatures (Sinyinda and Gramshaw, 1998; Kikuta and Erickson, 1968), these variation might be attributed to the diversity of the regional conditions (cultivar, environment and climate. etc) that might effect on the biosynthesis of compounds in different avocado fruits. The increasing prevalence of multi drug resistant strains with reduced susceptibility to antibiotics raises the specter of untreatable bacterial infections and adds urgency to the search for new infection fighting strategies (Sieradzki et al., 1999). Therefore, research into the effects of avocado oil is expected to enhance the use of this oil against diseases caused by the test pathogens, according to the clinical laboratory standard institution (CLSI, 2004). Antimicrobial screening of essential oil of avocado (Figure 3), was determined against *S.aureus, E. coli, P. aeruginosa and C. albicans* 100 µl of avocado oil at a dose of 100 mg/ml test by determination of the zone of inhibition and showed high inhibit activity against *P. aeruginosa* (18 mm), and moderate inhibit the activities against *S.aureus*, *C. albicans* and *E. coli* by 15, 14 and 13mm respectively, when compare with control; ampicillin for *S.aureus* by (21mm), Doxycycline by 25 mm and 24 mm for *E. coli* and *P. aeruginosa* respectively, and Nystatin for *C. albicans* by 23 mm. The lowest concentration of the sample required to inhibit the growth of test organism, was detected for each organism as MIC. The volatile oil was dissolved in dimethyl sulfoxide 0.2 ml DMSO/10 ml medium. MIC values are 6.4, 3.2, 3.2, 6.1 mg/ml for *S.aureus*, *E. coli, P. aeruginosa,* and *C. albicans* respectively (Table 2). From the previous antimicrobial study, the results obtained indicated the existence of antimicrobial compounds in avocado oil, therefore, its helpfulness in the management of many diseases that could be as a cause of infection (Guzman-Rodriguez et al., 2013). As presented in Table 3, avocado oil was able to reduce the blue DPPH-radical methanolic solution (125 µL of µM). Avocado oil was found to be half the potency of ascorbic acid at doses of 12.5, 25, 50, 100 µg/ml as well as it showed a significant antioxidant activity at a dose dependent matter of avocado oil by 70.95% and 71.85% at 200 and 400 µg\mL when compare to ascorbic acid and BHT, (Figure 4), respectively. The high antioxidant potency of oil was thus found to be partially correlated to its rich of alkanals and hydrocarbon content. On the bases of above finding, it was expected that the avocado oil would exhibit a considerable protective activity against the oxidative stress induced by free radicals. To verify these findings, avocado oil was subjected to the evaluation of their quorum sensing and biofilm formation of Avocado oil against pathogens. In the current study, sub-MIC concentrations (0.1-0.8 %v/v) of avocado oil were inhibited violacein production in wild-type *C. violaceum* CVO26 strain in concentration dependent action without affecting the population of the bacteria. The purple pigment (Violacein) production in *C. violaceum* is a QS regulated process, and its production is organized by CviIR-dependent QS system. Maximum reduction of 94% was recorded at 0.8 % v/v while at lower concentrations (0.1-0.4 % v/v) 45–80% decrease in violacein was noticed, (Figure 5). This dose-dependent manner of avocado oil on violacein production is in accordance with the reports on Indian medicinal herbs (Zahin et al., 2010), *Capparis spinosa* and *Cuminum cyminum* extracts (Packiavathy et al., 2012).

Virulence factors (VF) are known to play an important role during the invasion of the host cells. *P. aeruginosa* produces a range of QS-regulated VFs including elastase, protease and chitinase (Adonizio et al., 2008). This data corroborated with the literature where, total proteolytic chitinase and elastase activities of *P. aeruginosa* was decreased to varying levels by different plant extracts and volatile oils (Husain and Ahmad, 2013; Vattem et al., 2007).

Effect of avocado oil sub-inhibitory concentrations on virulence factors of *P. aeruginosa* PAO1 is shown in Figure 6a. Statistically significant decrease in **LasB** elastase activity was observed in the culture supernatant of PAO1 treated with sub-MICs of avocado oil. A minimum of 30% inhibition was observed when PAO1 was cultured with avocado oil at a concentration of 0.4 % (v/v) and maximum of 80% inhibition was observed at 3.2% (v/v) concentration of the oil. Elastase enzyme enhances the growth and invasiveness of the pathogen by degrading the structural components of the infected tissue (Kharazmi, 1989). In this current investigation, the avocado oil demonstrated concentration-dependent inhibition of elastase in PAO1, as shown in Figure 6a. This result is in alignment with the previous study (Musthafa et al., 2010) who demonstrated significantly inhibition of LasB activity by edible fruits. Early reports suggest that flavonoid rich extracts of edible plants exert an inhibitory effect against the QS dependent expression of proteolytic enzymes such as LasB in PA01. In addition, *Trigonella foenum-graceum* seed extract have been reported to inhibit elastase activity to certain levels (Husain et al., 2015).

Production of Pyocyanin (PP) is an important VF produced under QS regulation. The major role of PP is documented in pathogenesis mainly in cystic fibrosis (Winstanley, 2009). A similar reduction in PP production was reported in extracts of *Terminalia chebula*(Sarabhai, et al., 2013). A “Pyocyanin” has a green color produced by *P. aeruginosa* (PAOI) after 24-48 h of growth. The disappearance of this pigment indicated the lower levels of PP are found in the supernatant. The effect of avocado oil on the PP was performed. In the current study, PP level in *Pseudomonas* culture handled with this oil was significantly reduced without affecting the growth of bacteria against the green color of untreated cultures. This might be interpreted as quorum-control of PP(Dietrich, et al., 2006). Pyocyanin and its precursor phenazine-1-carboxylic acid (PCA) cause neutrophil apoptosis and impair neutrophil-mediated host defenses (Fothergill et al., 2007). Avocado oil at sub-lethal concentrations exhibited a considerable decrease in the PP by PAO1. The maximum significant reduction of 62% (*p* ≤0.005) in PP was recorded at a highest tested concentration (3.2%v/v) followed by 51, 38, and 15% in 1.6, 0.8, and 0.4 %v/v concentrations, respectively, Figure 6a. Our results are in agreement with the results of recent reports wherein Krishnan et al., (2012), and Gala et al., (2016) demonstrated that extracts of *Tinospora cordifolia* (stem) and *S. aromaticum* (bud) reduced PP significantly.

**Swarming motility and EPS** production by *P. aeruginosa* plays a pivotal role in the initiation, maturation, and maintenance of the biofilm architecture (Pratt and Kolter, 1998; Hentzer et al., 2003). So, any interference with the motility and exopolysaccharide production is bound to affect the BF by the pathogen. In the current study, treatment of PAO1 with sub-MICs of avocado oil showed significantly decrement of exopolysaccharide production, the extract (0.4–3.2 %v/v) demonstrated inhibition in exopolysaccharide production to the level of 19–72%. Similarly, swarming migration of PAO1 was also impaired considerably (10–69 %) after treatment with avocado oil (Figure 6a). This statistically significant reduction of motility and exopolymeric material is reported with *Trigonella foenum-graceum* seed extract(Husain et al., 2015). Elimination of *Pseudomonas* motility confirmed the potential effect of *C. olitorius* L. aqueous fraction of biofilm formation as modulation of bacterial motilities is associated with thinner and dispersed biofilm (Shrout et al., 2006).

**BF** is a drug resistant complex aggregation of microorganisms and is a key factor in the pathogenesis of *P. aeruginosa* (Caraher et al., 2007). In a biofilm adherent cells become embedded within a slimy extracellular matrix that is composed of extracellular polymeric substances (EPS). Thus, this oil indirectly demonstrated consequences on BF of all the target pathogens in part by interfering with its ability to reach the substratum and subsequent BF by disturbing AHL-mediated QS-system. It has also been proven that surface conditioning promotes surface adhesion and subsequent - microcolony formation (Sandasi et al., 2010). Biofilms are the cause of severe persistent infection and BF is considered as one of the potential drug targets to combat drug-resistant chronic infections (Hall-Stoodley et al., 2004; Wu et al., 2015). The avocado oil showed 32, 52, 72, and 83%, a significant decrease in the BF ability of PAO1 at 0.4, 0.8, 1.6, and 3.2 %v/v of oil concentrations, respectively, Figure 6b. Our observations find support from previous experimentation on BI in PAO1 by polyphenolic extract of South Florida plants (Adonizio et al., 2008) *Lagerstroemia speciosa* fruit extract (Singh et al., 2012), *Rosa rugose* (Zhang et al., 2014), standardized extract of *Sclerocarya birrea* (Sarkar et al., 2014), *Trigonella foenumgraceum seed* extract (Husain et al., 2015) and *Mangifera indica* leaf extract (Husain et al., 2017).

## CONCLUSIONS

Avocado oil is known for its medicinal use and our study appends an additional note on its QS and BI properties against pathogenic bacteria. The current study demonstrates that avocado oil could inhibit the QS mediated virulence factor production in *C. violaceum* and *P. aeruginosa*. Moreover, the treatment with sub-MICs of avocado oil significantly inhibited the QS-mediated BF, EPS production and swarming motility in these pathogens. Wide-spectrum *in vitro* inhibition of QS controlled virulence factors such as violacein, elastase, Pyocyanin, EPS and biofilm in test pathogens was determined. Thus, these results postulated that avocado oil has powerful anti-infective properties and could confirm to be an effective anti-QS and antibiofilm agent against pathogens.

## MATERIALS AND METHODS

### Materials

Commercial avocado oil was purchased from a local market in Riyadh, Saudi Arabia under the trade name of Yasin, 100% natural and unrefined avocado oil were acquired from a Company labeled Nobel Foods SA.de CV, Mexico. Physically, avocado oil is a dark yellow liquid with a characteristic aromatic odor, soluble in ether, chloroform and insoluble in water.

#### Microorganisms (MOs)

American type of culture collection (ATCC) standard against various microorganisms namely, *Staphylococcus aureus* (ATCC25922), *Escherichia coli* (ATCC25923), *Pseudomonas aeruginosa* (ATCCPAO1) and *Candida albicans* (SC315) were used to investigate the antibacterial activity of avocado oil.

### Methods

#### Bacterial strains, media and growth conditions

The bacterial strains which were used in this study were *C. violaceum* CV026 (a mini-Tn5 mutant of *C. violaceum* 31532 that cannot synthesize its own AHL, but responds to exogenous C4 and C6 AHLs) and *P. aeruginosa* PAO1 (C4 and 3-oxo-C12 HSL producer (McLean et al., 2004). Luria-Bertani (LB) medium was used to grow the bacterial strains at 30 ºC for 24 h. However, *C. violaceum* CV026 medium with hexanoyl homoserine lactone was supplied by (C6-HSL; Sigma-Aldrich, St Louis, MO, USA).

#### Gas chromatography/Mass spectrometry (GC/MS)

The GC-MS analysis was performed in a Perkin Elmer Clarus 600 gas chromatograph inked to a mass spectrometer (Turbo Mass) available at Central Laboratory, College of Pharmacy, King Saud University, Riyadh. An aliquot of 1 µL of the extract was injected into the GC column Elite -5 MS of 30 m long, 0.25 µm film thickness, 0.25 mm internal diameters.

#### Capillary column using the following temperature program

The GC-MS system starts with the initial oven temperature of 40 ºC increasing at a rate of 5 ºC/min, and then oven final first ramp 100 ºC at a rate of 5 ºC for 2 minutes, the oven ramp rate 5 ºC/min transfer line heater 200 ºC, oven final temperature 300 ºC for 5 minutes. The injector temperature was maintained at 220 ºC. The inlet temperature was 300 ºC MSD solvent delay 3.5 minutes. Helium was used as a mobile phase at a flow rate of 1.0 mL/min. Mass spectral detection was carried out in the electron ionization mode by scanning at 40 to 600 a.m.u. Finally, unknown compound was identified by comparing the spectra with that of the National Institute of Standard and Technology library (NIST 2005) and Wiley Library 2006 (Ver 2.1). The total time required for analyzing a single sample was 61 minutes. The Retention Index (RI) was calculated by running the standard solution of C-7 to C-30 saturated alkane’s standard from SUPELCO with the same method as a sample. The concentration of alkanes was 1000 µg/ml. The RI values were calculated by AMDIS software 32.

#### Identification of Components by GC-MS

GC retention time was used to identify the components and matching them with Wiley, 2006 library as well as by comparison the fragmentation patterns of their mass spectra with those reported in the literatures (Adams, 1995; Mclafferty and Staffer, 1989) and several identified components were identified as sterols, fatty acids, alkanes and alcohols compounds. A total of 23 detectable peaks was selected from avocado oil.

#### Antibacterial Assay

The agar well diffusion method Perez *et al*., (1990) as adopted earlier Ahmad and Beg, (2001). Briefly, Sabouraud Dextrose (SD) and Soyabean Casien Digest (SCD) were used for *S. aureus, E. coli, P. aeruginosa and C. albicans* test bacteria, respectively. Freshly prepared microbial cultures were appropriately grown at 37°C in sterile normal saline solution to obtain the cell suspension at 105 CFU: mL

#### Determination of minimum inhibitory concentration (MIC) of avocado oil

Minimum inhibitory concentration of avocado oil against drug resistant clinical strains was determined by a broth dilution method, using specific dye (p-iodonitro tetrazolium violet) as an indicator of growth as described by Eloff, (1998). ^[53]^ Briefly, 2 mL of Muller-Hinton broth was mixed with 2 mL of avocado oils and were serially diluted. 2 mL of different actively grown culture of test strains was added before incubating for overnight, at 37°C. 0.8 mL of 0.02 mg/mL of indicator dye (p-iodonitro tetrazolium violet) was added to each tube after examining turbidity visually, and incubated at 37°C. The color development of each tube was examined after 30 min. Absence of growth was also confirmed, with the addition of 0.1 mL of broth from each test tube to normal nutrient agar plates. MIC is defined as the minimum concentration of avocado oil, which inhibited the visible growth of the test strains.

#### Biosensor bioassay detection of anti-quorum sensing activity

The anti-QS activity of avocado oil was detected by bioassay using the reporter strains *C. violaceum* CV026 and *P. aeruginosa* PAO1. To carry out this study, various concentrations of avocado oil ranges of 0.4-0.8 and 0.4-3.2 % (v/v) were loaded onto 6-mm sterile discs and placed on the surface of C. violaceum CV026 and *P. aeruginosa* PAO1, respectively. Then LB agar plates supplemented with 50 mL 1 mg mL-1 C6-HSL were incubated for 24-48h. The negative control used for this test was discs loaded with ethanol. A zone of colorless, but viable cells around the disc revealed the QS inhibition.

#### Quantitative estimation of violacein

Extent of violacein production by *C. violaceum* (CVO26) in presence of Sub-MICs of avocado oil was studied by extracting violacein and quantifying photometrically using the method of Blosser and Gray, (2000) with little modifications (Husain et al., 2015). About 10 mL LB broth containing different concentrations of avocado oil was inoculated with 100 mL C. violaceum ATCC12472 (106 CFU/mL). Similarly, control solvent was prepared and all the tubes with continuous orbital shaking at 130 rpm were incubated for 24 hours. Water soluble Violacein was extracted with n-butanol from the cells and was spectrophotometrically quantified at an optical density (OD) 585 (UV-1800; Shimadzu). In order to find the effect of oil on bacterial growth, serial dilution of culture grown in the presence of oil was measured by the standard plate count method.

#### Effect on virulence factor production

The effect of Sub-MICs of test agents on virulence factors of *P. aeruginosa* such as LasB elastase, pyocyanin, swarming motility, EPS extraction and quantification was assessed as described previously (Husain et al., 2013). The avocado oil’s effect on pyocyanin pigment production in *P. aeruginosa* PAO1 was determined by growing *P. aeruginosa* PAO1 in glycerol alanine minimal medium supplemented with different concentrations of avocado oil and incubated for 24 hours. Chloroform was used to extract pyocyanin from the cell-free supernatant and acidified with 0.2 M HCl, which was spectrophotometrically by recording on OD520.

Luria broth (LB) semisolid (0.5% agar) medium supplemented with avocado oil was used to perform swarming assay. The swarming diameter was measured after 24 h incubation of *P. aeruginosa* PAO1 with inoculated LB agar plates (Vattem et al., 2007) Elastolytic and proteolytic inhibition activities was assessed by gowing *P. aeruginosa* PAO1 in LB medium supplemented with different concentrations of avocado oil and incubated for 16 hours. To 900 mL elastin congo red (ECR) buffer (100 mM Tris, 1 mM CaCl2, pH 7.5), 100 mL of culture supernatant was added which was containing 20 mg of ECR (Sigma-Aldrich, Hamburge, Germany) and incubated at 37º C for 3 hours (Kessler et al., 1982). After removing the insoluble ECR by centrifugation, the absorbance of the supernatant was spectrophotometrically measured at OD495. To 900 mL ECR buffer containing 3 mg azocasein (Sigma-Aldrich, Hamburge, Germany), 100 mL culture supernatant was added and incubated at 37ºC for 30 minutes. To each reaction tube 100 mL of trichloroacetic acid (10 %) was added. After 30 minutes, the tubes were centrifuged and absorbance of the supernatant was determined by reading OD440.

#### Assay for biofilm inhibition

O’Toole and Kotler (1998) microtitre plate assay method was performed to determine the effect of avocado oil for BF formation. To 1 mL of fresh LB medium, 1% overnight cultures (0.4 OD at 600 nm) of test pathogens were added in the absence and presence of sub-MICs of test agents. Bacteria were allowed to adhere and grow without agitation for 24 h at 30°C. Media along with free-floating planktonic cells were removed from microtitre plate after incubation and rinsed twice with sterile water. The formed BF was stained with 0.1 % crystal violet (200 μL) solution. After 20 min, crystal violet was completely drained and 200 μL of 95% ethanol was added to the wells to solubilize the crystal violet from the stained cells. Then the microplate reader was used to quantify the BF biomass by measuring the absorbance of BF biomass at OD 470 nm.

#### Antioxidant assay (DPPH radical scavenging assay)

The free radical scavenging activity of avocado oil against stable 1,1-diphenyl-2-picrylhydrazyl (DPPH) was determined spectrophotometrically by slightly modified method of Gyamfi *et al*. (1999) Different concentrations of avocado oil were mixed with 150 mL DPPH to obtain the final concentration of 100 mM. The reaction was incubated in the dark for 30 min at 37° C. At 515 nm, optical density was measured. Antioxidant activity was expressed as IC50. Ascorbic acid was used as standard antioxidants for comparison. Methanol was used as a control. Quadruplicates were used to determine all curves.

#### Determination of minimum inhibitory concentration

MIC of the avocado oil was determined after incubation at 37ºC for 18 hrs, against selected pathogens using broth macrodilution method.(CLSI, 2004) Sub-MICs were selected for the assessment of anti-virulence and anti-biofilm activity in the above test strains.

## Statistical analysis

All experiments were performed in triplicates and the data obtained from experiments were presented as mean values and the difference between control and test were analyzed using student’s *t* test.

## Author Contributions: Authors’ contributions

HA-Y: Designed the study suggested; MA: Performed the experiment and interpreted the data; MF: Write the manuscript and data collection; SR: perform GC/MS analysis; WH: helped in literature survey.

## Acknowledgment

This research project was supported by a grant from the “Research Center of the Center for Female Scientific and Medical Colleges”, Deanship of Scientific Research, King Saud University

